# Delayed growth of SK-ES-1 Ewing sarcoma tumors associated with reduced expression and activity of Trk receptors, PI3K, and IGF1R

**DOI:** 10.1101/2025.10.23.684233

**Authors:** Bruna Almeida dos Santos, Natália Hogetop Freire, Livia Fratini, Matheus Dalmolin, Josimar Macedo de Castro, Marcelo A. C. Fernandes, Lauro Gregianin, André T. Brunetto, Algemir Lunardi Brunetto, Luis Fernando da Rosa Rivero, Mariane da Cunha Jaeger, Rafael Roesler, Caroline Brunetto de Farias

## Abstract

Ewing sarcoma (ES) is an aggressive childhood tumor. We have previously shown that tropomyosin receptor kinase (Trk) neurotrophin receptors regulate viability, and the pan-Trk and kinase inhibitor K252a restores sensitivity to chemotherapy, in ES cells. Here, we show that administration of K252a in mice delays ES tumor growth and reduces expression and activity of multiple targets. BALB/c nu/nu nude mice inoculated with SK-ES-1 ES cells were given daily intraperitoneal (i.p.) injections of K252a for 18 days. Immunohistochemical analysis of tumors was performed to quantify the expression and phosphorylation of Trk receptors, phosphoinositide 3-kinase (PI3K), and insulin-like growth factor 1 receptor (IGF1R). Associations between genes encoding Trks, PI3G, and IGF1R and overall survival (OS) in patients with ES was examined. Treatment with K252a led to a transient delay in ES tumor growth and reduced the expression and phosphorylation of TrkA, TrkB, PI3K, and IGF1R. Combined treatment with K252a and the IGF1R inhibitor NVP-ADW742 was more effective in reducing ES cell viability than each compound alone. Significant associations between expression of *NTRK* genes and patient OS were found, indicating that *NTRK* genes should be further evaluated as biomarkers for prognosis in patients with ES.

## INTRODUCTION

Ewing sarcoma (ES) is an aggressive type of childhood cancer that can arise either in bone or soft tissues, with a peak incidence during adolescence, though it also affects younger children and can occur at any age. ES tumors exhibit undifferentiated features that reflect elements of both mesenchymal and neural lineages. Although their exact cellular origin remains debated, current evidence suggests that ES may arise from neural crest–derived or mesenchymal stem cells, and that disruptions in epigenetic programming and multiple signaling pathways during embryonic development can contribute to tumor initiation and progression. At the genetic level, Ewing sarcoma is the prototypical example of a cancer driven by a specific chromosomal translocation. The translocations involve members of the FET (FUS, EWSR1, TAF15) and ETS transcription factor families, most commonly generating the EWS::FLI1 fusion oncogene. The resulting aberrant transcription factor reprograms gene expression and profoundly alters cell fate and differentiation potential, conferring the tumor’s hallmark biological behavior. Therapeutically, the introduction of multimodal cytotoxic regimens has markedly improved outcomes for patients with localized disease, yielding survival rates of approximately 70–80%. However, prognosis remains poor for those with metastatic or relapsed ES, with long-term survival dropping to around 30%. Moreover, survivors frequently endure substantial, long-lasting adverse effects from treatment, including growth impairment, organ toxicity, and limb amputations, alongside the emotional and social challenges that come with surviving a severe childhood cancer [1–6].

Given that the oncogenic fusion protein EWS::FLI1 by itself is hard to directly target with drugs, other components of the ES cell signaling landscape have been investigated as therapeutic targets. These include cell surface receptors such as receptor tyrosine kinases (RTKs) [7–10]. Members of the RTK subfamily, tropomyosin receptor kinase (Trk) receptors include TrkA, TrkB, and TrkC, which are encoded by the NTRK1, NTRK2, and NTRK3 genes, respectively. TrkA is the high-affinity receptor for nerve growth factor (NGF), while TrkB primarily binds brain-derived neurotrophic factor (BDNF) and neurotrophin-4 (NT-4). Neurotrophin-3 (NT-3) has the highest affinity for TrkC but can also to a lesser degree activate TrkA and TrkB. Upon ligand binding, Trk receptors undergo homodimerization and autophosphorylation at specific tyrosine residues, stimulating downstream cell signaling cascades that include the phosphoinositide 3-kinase (PI3K), mitogen-activated protein kinase (MAPK), and phospholipase C-γ (PLCγ)/protein kinase C (PKC) pathways [11–12]. Beyond their physiological roles in the nervous system, Trk receptors have been increasingly implicated in tumor progression, therapy resistance, and cell survival in several types of cancer [13–17]. Trk inhibitors larotrectinib and entrectinib are currently used to treat patients with tumors harboring NTRK gene fusions, regardless of cancer type [13, 18–21].

In sarcomas, NTRK fusion genes were first identified in infantile fibrosarcoma (IFS), which can harbor a canonical ETV6-NTRK3 gene fusion [22, 23]. Current evidence demonstrates various NTRK1 and NTRK3 fusion genes in IFS and other sarcoma types [23, 24]. TRK fusion proteins show activation of the kinase domain, which can lead to constitutive activation of PI3K and MAPK signaling through a mechanism dependent on insulin-like growth factor 1 receptors (IGF1R) [23, 25]. Patients with NTRK fusion-positive IFS tumors show a 96% response rate to larotrectinib, indicating that NTRK fusions define a molecular subset of sarcoma tumors highly sensitive to Trk inhibitors [26].

In a previous study, we reported that TrkA and TrkB are expressed in ES tumors, and treating ES cells to selective TrkA or TrkB inhibitors reduced cell proliferation, an effect that was increased when TrkA and TrkB were blocked simultaneously [27]. We then used the furanosylated indolocarbazole K252a, a microbial alkaloid originally isolated from the soil fungi *Nocardiopisis sp.* and structurally related to staurosporine, as a pan-Trk inhibitor [28–31]. Treatment with K252a induced changes in cell morphology, reduced β-III tubulin levels, and decreased mRNA expression of NGF, BDNF, TrkA and TrkB. In addition, combining K252a with subeffective doses of cytotoxic chemotherapeutic drugs resulted in decreased ES cell proliferation and survival and restored sensitivity to chemotherapy in chemoresistant ES cells [27]. Here, we show that treatment with K252a delays ES tumor growth in mice and the effect is associated with inhibition of multiple tumor targets.

## RESULTS

### Systemic administration of K252a produces a transient delay in ES tumor growth

Treating nude mice inoculated with human SK-ES-1 ES cells with daily injections of K252a resulted in a significant reduction in tumor growth during treatment compared to controls. There were significant effects of treatment and time as well significant interactions between these factors (all *p*s < 0.01; Figure 1B, 1C). The halt in tumor growth was apparent between treatment days 9 and 15. However, by day 18, tumor size in the treatment group had recovered to levels comparable to those observed in the control group, suggesting that the drug effect was transient (Figure 1B). There were no significant differences between groups in body weight (Figure 1D) or serum biochemical markers, although apparent reductions in AST and ALT levels were observed in K252a-treated mice (Figure 1E-H).

**Figure 1:**
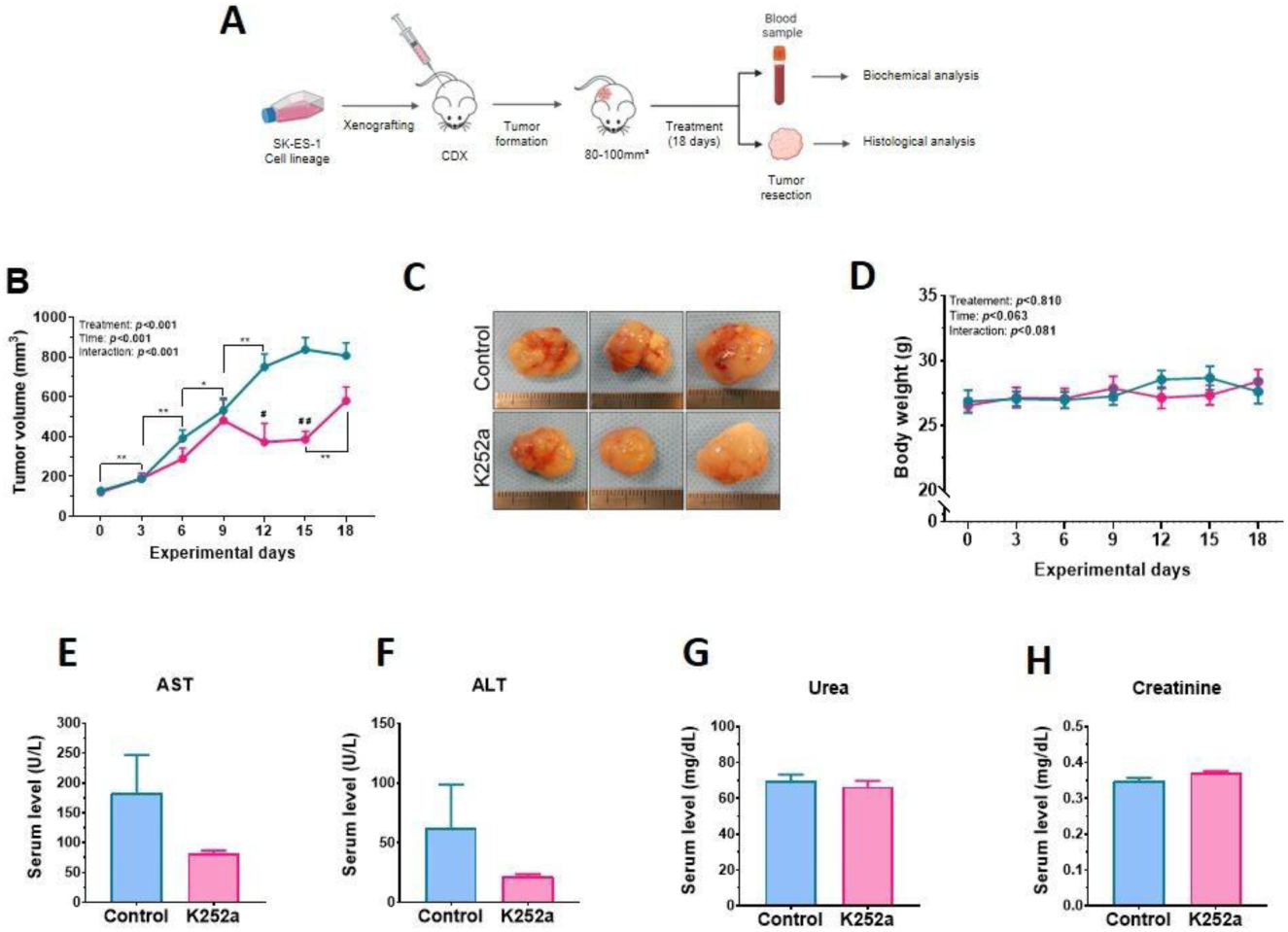
Treatment with K252a delays ES tumor growth. (**A**) Schematic of the experimental design. Human SK-ES-1 ES cells were xenografted into nude mice and the tumors were allowed to grow for 18 days. (**B**) tumor volume across experimental days. The blue line represents control mice and the pink line represents mice given K252a. The number of alive animals in each experimental day was: day 0, control, *n* = 14; K252a, *n* = 13; day 3, control, *n* = 14; K252a, *n* = 13; day 6, control, *n* = 14; K252a, *n* = 13; day 9, control, *n* = 14; K252a, *n* = 13; day 12, control, *n* = 14; K252a, *n* = 12; day 15, control, *n* = 11; K252a, *n* = 9; day 18, control, *n* = 7; K252a, *n* = 7; * *p* < 0.05, ** *p* < 0.01, comparisons between experimental days in each group. (**C**) Representative tumors from controls and K252-treated mice. (**D**) Body weight across experimental days; Serum levels of (**E**) AST, (**F**) ALT, (**G**) urea, and (**H**) creatinine in controls and K252a-treated mice (*n* = 7 per group).

### Treatment with K252a reduces the protein expression and phosphorylation of TrkA, TrkB, PI3K, and IGF1R, without affecting TrkC in ES tumors

Tumors from mice treated with K252a had reduced protein levels of total (Figure 2) and phosphorylated (Figure 3) TrkA (total, *p* = 0.06, phosphorylated, *p* = 0.04), TrkB (total, *p* = 0.05, phosphorylated, *p* = 0.012), and PI3K (total, *p* = 0.05, phosphorylated, *p* = 0.02), but unchanged TrkC levels. A subsequent analysis showed that K252a also reduced total and phosphorylated levels of IGF1R (total, *p* = 0.02, phosphorylated, *p* = 0.01; Figure 4A, 4B). Significant correlations were found in reduction of protein expression between several pairs of targets in cohorts of ES tumors from both groups of mice (Supplementary Figure S1).

**Figure 2:**
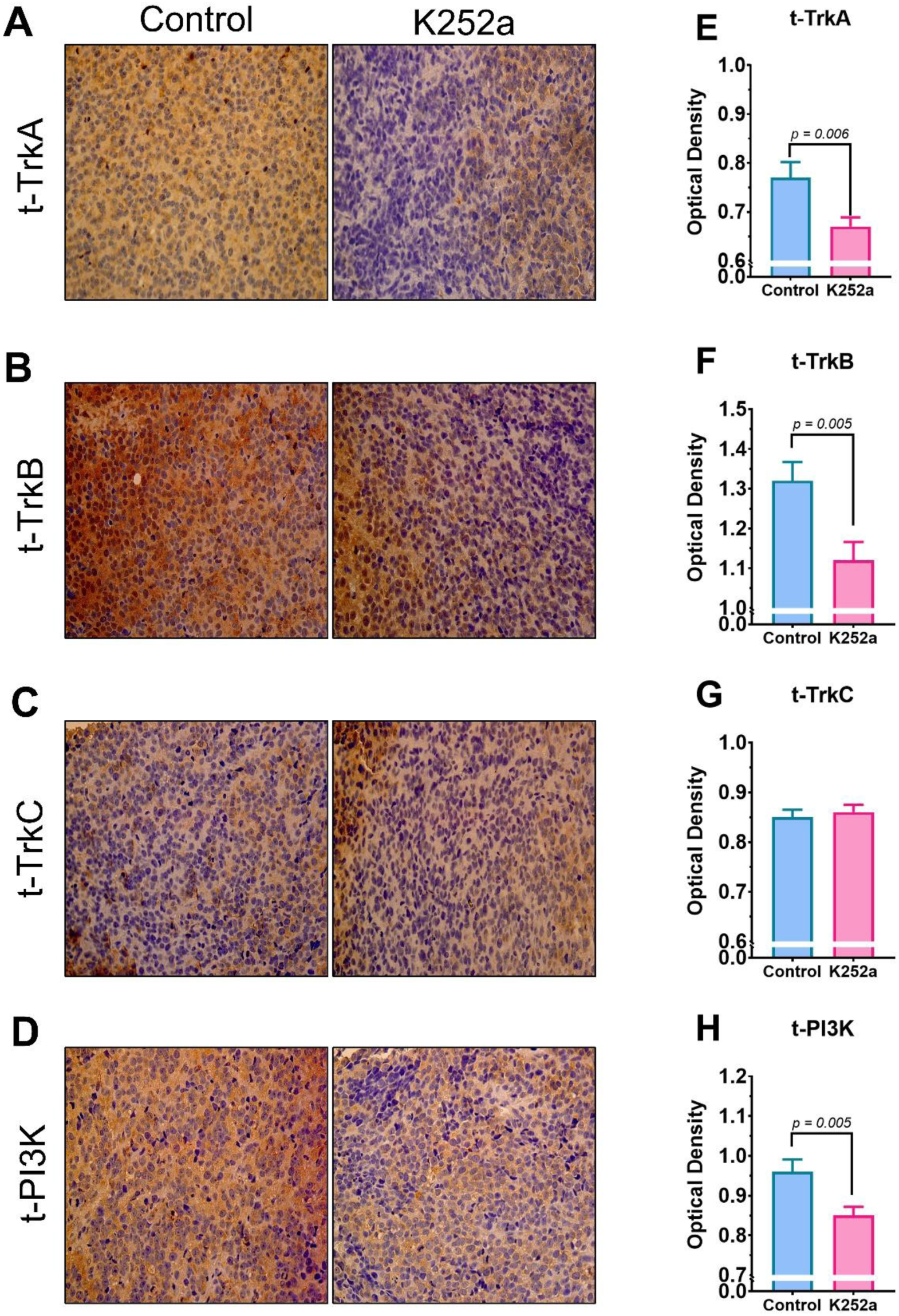
Immunohistochemical expression of total (phosphorylated and non-phosphorylated, t) (A, E) TrkA, (B, F) TrkB, (C, G) TrkC, and (D, H) PI3K in ES tumors from controls and K252a-treated mice (n = 7 per group). (**A**-**D**) Representative micrographs, X 40. (**E**-**H**) Protein content was quantified and shown as mean + S.E.M. optical density; *p* values are indicated in the panels.

**Figure 3:**
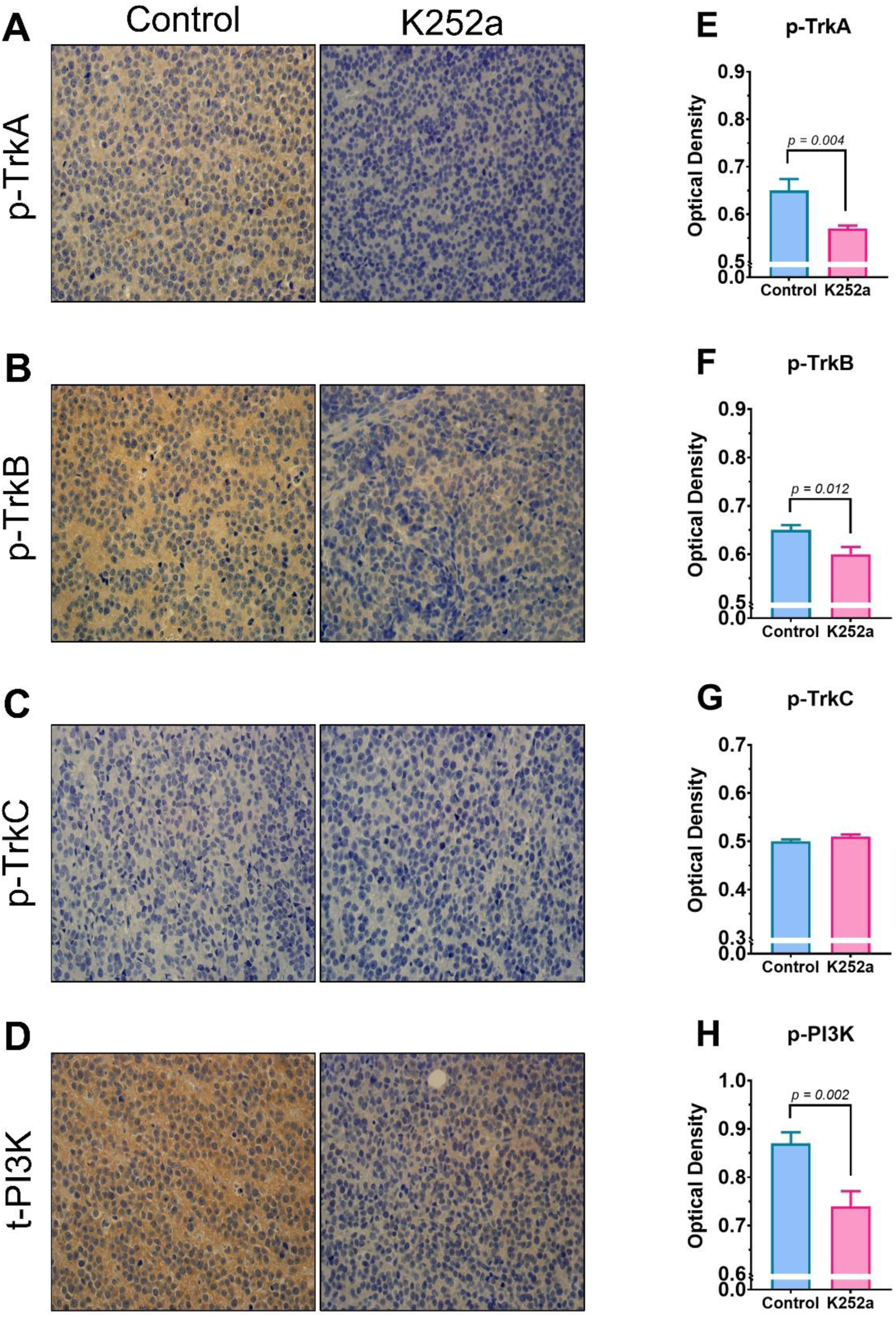
Immunohistochemical expression of phosphorylated (p) (A, E) TrkA, (B, F) TrkB, (C, G) TrkC, and (D, H) PI3K in ES tumors from controls and K252a-treated mice (*n* = 7 per group). (**A**-**D**) Representative micrographs, X 40. (**E**-**H**) Protein content was quantified and shown as mean + S.E.M. optical density; *p* values are indicated in the panels.

**Figure 4:**
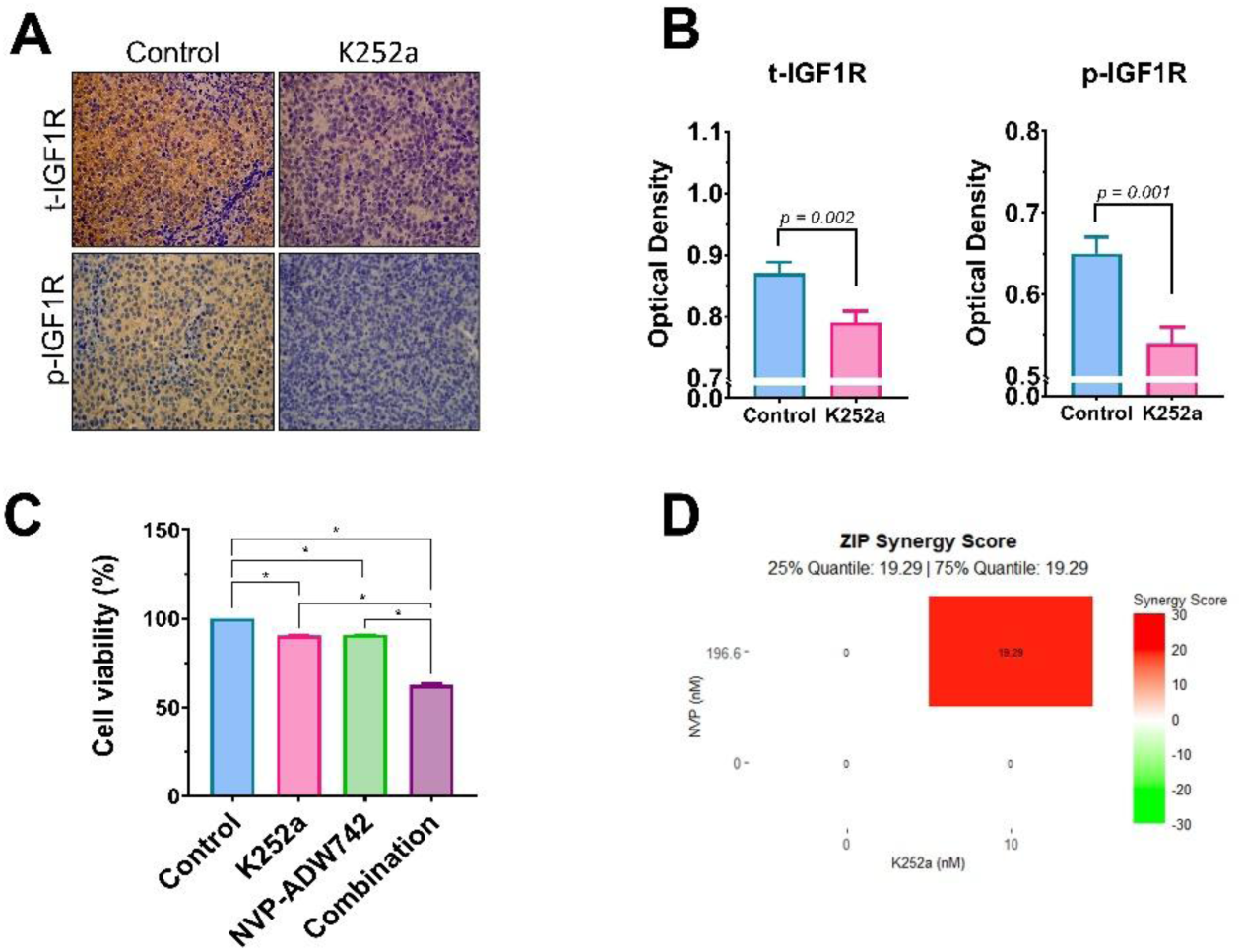
IGF1R expression and interaction with K252a in ES tumors. **(A) Representative micrographs (X 40) of immunohistochemical expression for total (t-IGF1R) and phosphorylated (p-IGF1R) IGF1R in controls and K252a-treated mice (*n* = 7 per group).** (**B**) Protein content was quantified and shown as mean + S.E.M. optical density; *p* values are indicated in the panels. (**C**) Viability of SK-ES-1 ES cells treated with K252a (10 nM), the IGF1R inhibitor NVP-ADW742 (196.6 nM), or the two inhibitors combined. Cell viability was assessed by a trypan blue cell counting assay; *n* = 3 independent experiments, *p* < 0.05. (**D**) ZIP synergism score for the combination of K252a and NVP-ADW742.

### Combination treatment with K252a and an IGF1R inhibitor reduces ES cell viability

Exposure to K252a or NVP-ADW742 at doses equivalent to the IC10 resulted in a small but statistically significant reduction in the viability of SK-ES-1 ES cells. When the two drugs were used in combination, the resulting inhibition of cell viability was significantly more pronounced compared to the effects of each drug given alone (all *p*s < 0.05, Figure 4C). Significant synergism was demonstrated between the two drugs (25 and 75% quantiles = 19.29, Figure 4D).

### NTRK gene expression is associated with OS in patients with ES

TrkB, encoded by the NTRK2 gene, is considered a preferred target of K252a inhibition. We found that higher NTRK2 transcription is associated with worse prognosis assessed by shorter OS, specifically in the COG cohort (*p* = 0.0042, Figure 5C). In contrast, high NTRK1 expression was associated with longer OS in the EuroEwing cohort (*p* = 0.0049, Figure 5B). NTRK3 showed opposite patterns of association in the two datasets, indicating shorter OS in the COG cohort (*p* = 0.0052, Figure 5E) but longer OS in the EuroEwing cohort (*p* = 0.013, Figure 5F). No significant associations between gene expression and OS were found for PIK3CA (Supplementary Figure S2A, S2B), PIK3R1 (Supplementary Figure S2C, S2D), or IGF1R (Supplementary Figure S3).

**Figure 5:**
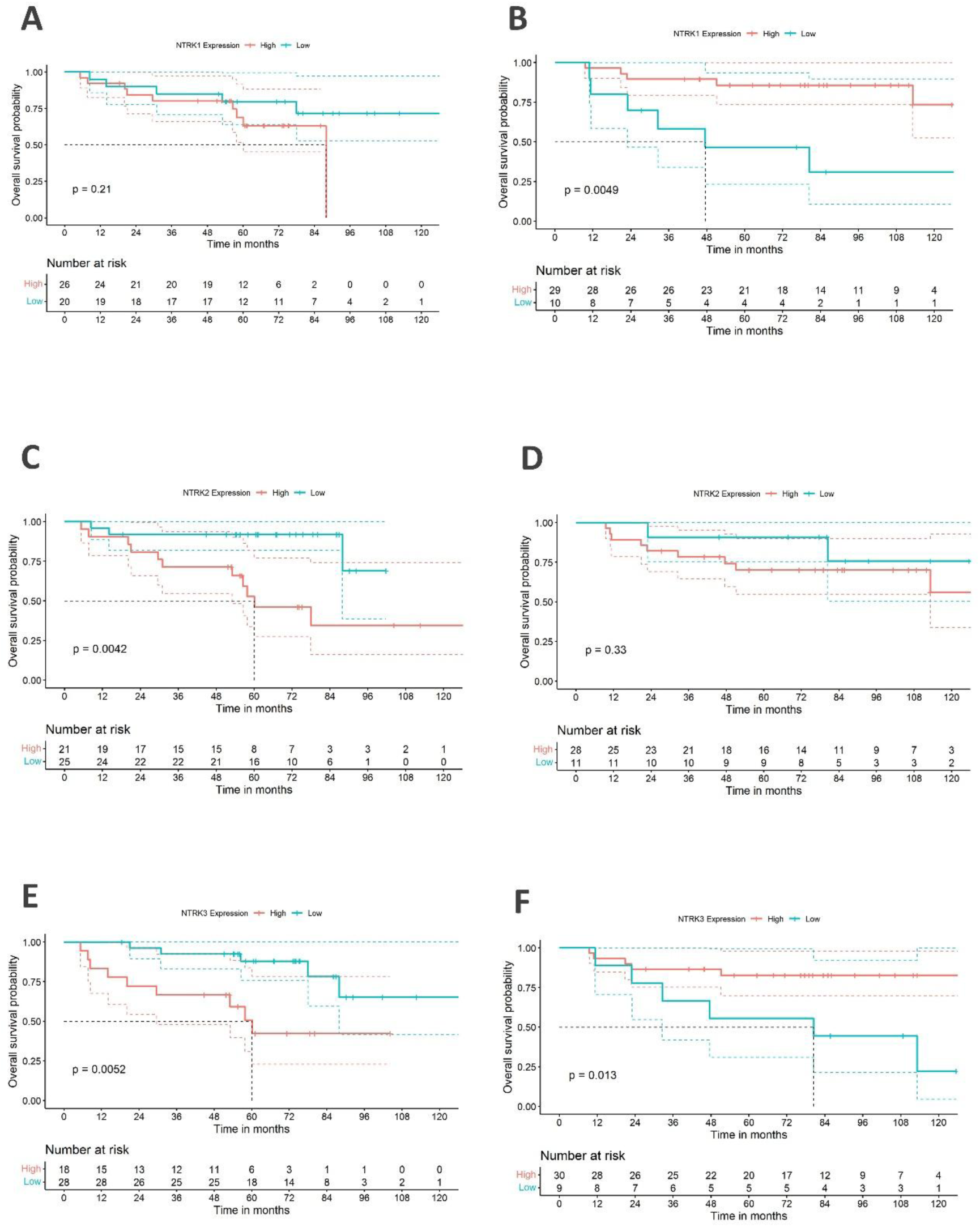
Associations between NTRK gene expression and prognosis assessed by OS in patients with ES. Datasets GSE63155 and GSE63156, comprising expression data from 46 primary ES tumors biopsies from COG and 39 tumors from the EuroEwing collaborative group respectively [58] were used (**A**) NTRK1, COG; (**B**) NTRK1, EuroEwing, (**C**) NTRK2, COG; (**D**) NTRK2, EuroEwing, (**E**) NTRK3, COG; (**F**) NTRK3, EuroEwing; *p* values are indicated in each panel.

## DISCUSSION

K252a has been widely used as a tool to investigate Trk-mediated signaling, particularly for TrkB, due to its potent inhibitory effects on Trks at nanomolar concentrations. However, at higher or even similar concentrations, and certainly at the doses we used in our *in vivo* experiment, it can also directly inhibit several other protein kinases [32–36]. Therefore, it is more accurate to interpret our findings as reflecting the effects of a non-specific, multi-kinase inhibitor, rather than those of a TrkB or pan-Trk inhibitor. Administration of K252a was able to significantly reduce the protein content and phosphorylation (a biochemical surrogate for kinase activation) of TrkA, TrkB, PI3K, and IGF1R in SK-ES-1 ES tumor xenografts, raising the possibility that either direct or indirect inhibition of these targets at least partially mediates the effect of K252a on tumor growth. Our correlation analysis showed that the expression of p-PI3K was positively associated with the content of both p-TrkA and p-TrkB, supporting the possibility that PI3K is among the downstream targets of Trk inhibition. Some of the main findings from the animal and cellular experiments reported in this study are summarized in Figure 6.

**Figure 6:**
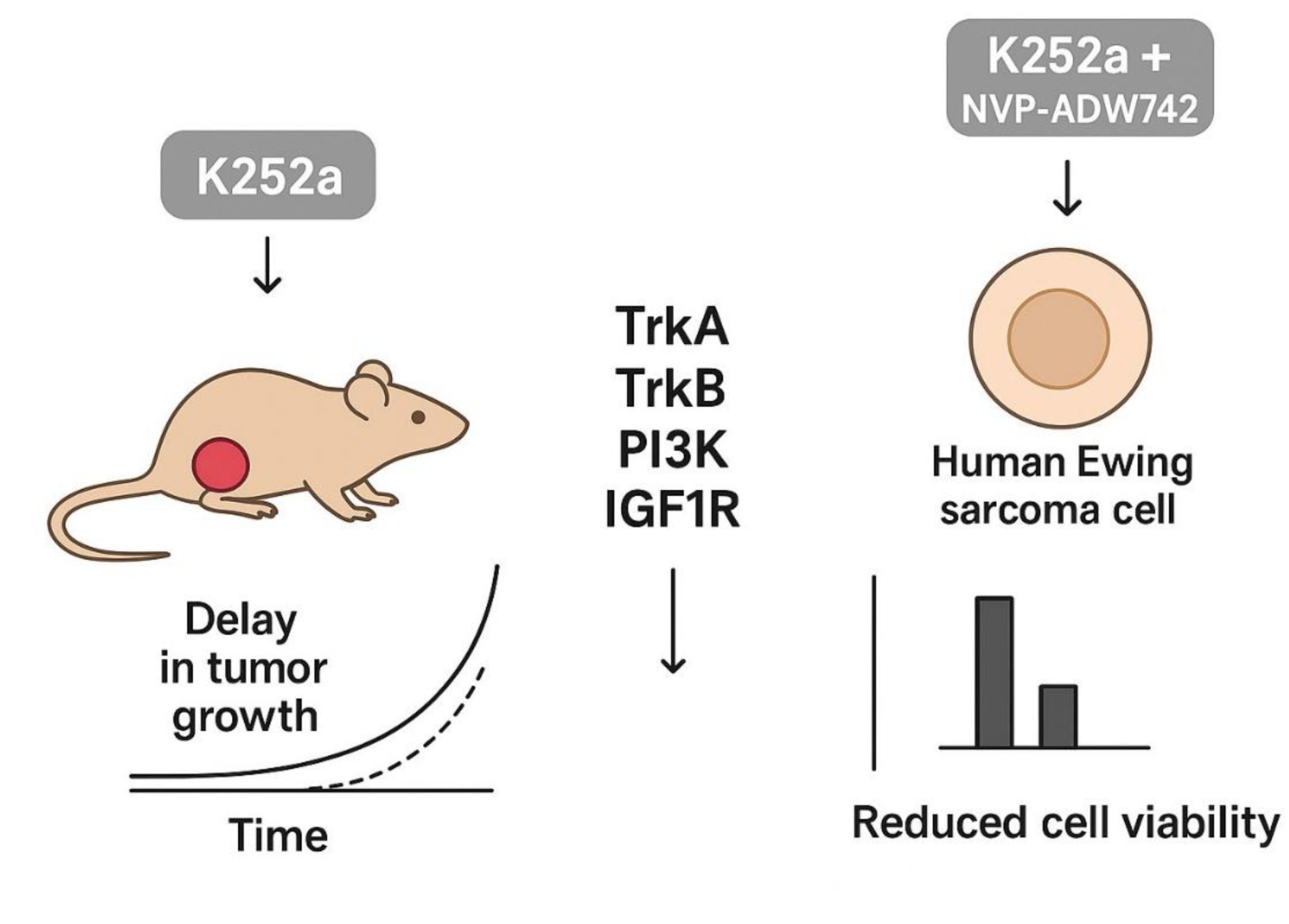
Schematic model highlighting selected aspects of K252a effects on ES. Systemic administration of K252a delays the growth of human SK-ES-1 ES tumors xenografted into nude mice and the effect is associated with reduced expression and phosphorylation of TrkA, TrkB, PI3K, and IGF1R in the tumors. Combined exposure to K252a and the IGF1R inhibitor NVP-ADW742 reduces SK-ES-1 ES cell viability more effectively than each compound alone.

One critical aspect of our findings was the transient nature of the antitumor effect of K252a. Tumors can rapidly acquire resistance to other Trk inhibitors through recurrent mutations in the Trk kinase domain [37]. However, it is more likely that rapid acquisition of resistance occurred through biochemical compensatory changes, rather than genetic alterations which can take longer to manifest. It has been shown that neuroblastoma cells overexpressing TrkB can developed resistance to entrectinib, a pan-TRK, anaplastic lymphoma kinase (ALK), and proto-oncogene tyrosine-protein kinase ROS (ROS1) inhibitor, by adaptations including activation of MAPK, downregulation of PTEN, and IGF1R activation. These changes allow preservation of downstream TrkB signaling in the presence of entrectinib [38]. It is also possible that the treatment rapidly selected for pre-existing resistant cells within the tumor.

Consistent with the possibility, raised by our previous study [27], that TrkA and TrkB expression and signaling influence ES growth, we found that higher NTRK2 gene expression can be associated with worse prognosis assessed by OS, whereas high NTRK1 expression indicates longer OS. These effects were observed specifically in one of the two cohorts examined, so that significant effects were found for NTRK1 in the EuroEwing dataset and for NTRK2 in the COG dataset. Contrasting patterns between the two patient cohorts were found for NTRK3; high gene expression was associated with a shorter OS in the COG dataset but longer OS in the EuroEwing dataset. We previously found a similar association between high NTRK3 transcription and longer survival in patients with medulloblastoma, the main type of pediatric brain tumor [17]. Both NTRK1 and NTRK3 can act either as oncogenes or tumor suppressor genes. NTRK1 is a tumor suppressor inhibited by methylation in ovarian cancer and endometrial hyperplasia [39, 40]. Reconstitution of normal NTRK3 expression previously suppressed by aberrant hypermethylation induces apoptosis in colon tumors [41], and NTRK3 is a tumor suppressor gene inactivated by methylation also in cervical cancer [42]. However, the first study reporting a role for the NTRK3 product, TrkC, in ES found the receptor to be frequently overexpressed in human metastatic ES cells, promoting cell survival, tumorigenicity, and metastasis, and loss of TrkC expression reduced experimental ES tumor growth and metastasis in mice [43]. Although NTRK gene fusions are mutually exclusive with the translocations that define ES [44], our previous *in vitro* findings and current results support the possibility that Trk inhibition can reduce ES growth even in the absence of pathological gene fusions.

Overexpression of IGF1R has been associated with more aggressive tumors and worse outcomes in ES, although our own transcriptional analysis did not reveal an association between IGF1R mRNA expression and survival of ES patients. Monoclonal antibodies targeting IGF1R have been developed and tested clinically in patients with ES in several trials, although the clinical efficacy remains to be fully established [45–50]. IGF1R activation enhances the expression of TrkA and TrkB at the mRNA and protein levels [51], and IGF1R functionally interacts with Trks in regulating drug resistance [25, 38]. Another novel finding reported in the present study is that the combination of K252a with an IGF1R inhibitor was more efficient in reducing ES cell viability than each drug given alone, highlighting the importance of evaluating combinatorial therapeutic regimens involving IGF1R inhibitors and drugs targeting Trks and protein kinases [52]. SK-ES-1 cells are derived from an ES tumor occurring in the bone of a 18 year-old male patient [53]. We describe the successful growth of ES tumors using SK-ES-1 cells xenografted into nude mice. Previous studies have used and validated SK-ES-1 xenografts as a preclinical model of ES [54, 55]. However, one of the limitations of our study is that patient-derived xenograft (PDX) mouse models, defined as models developed from direct implantation of patient tumor tissue into immunodeficient mice without prior cell culture or expansion, are now considered the gold standard for *in vivo* preclinical drug testing, and a number of such models have been developed for ES and other pediatric solid tumors [56]. It is also worth noting that our experiments do not allow to determine whether the reductions in target expression and phosphorylation are due to direct inhibition by K252a or downstream to drug-induced inhibition of Trks or other kinases. Future experiments should explore in detail the effects of selective TrkA, TrkB, and PI3K inhibitors, and compare them with multi-kinase inhibitors.

In summary, we report evidence suggesting that the combined inhibition of Trks, PI3K, and IGF1R by a multi-kinase inhibitor can delay ES tumor growth. In addition, NTRK genes should be further evaluated as potential biomarkers for prognosis in patients with ES.

## MATERIALS AND METHODS

### Ethics statement

All experimental procedures with animals were conducted in accordance with the guidelines established by the National Council for Animal Control and Experimentation (CONCEA, Ministry of Science, Technology and Innovation, Brazil), as well as with Animal Research: Reporting of In Vivo Experiments (ARRIVE) guidelines. All protocols used in this study were approved by the institutional Animal Care and Use Committee (CEUA, Hospital de Clínicas de Porto Alegre-HCPA), under project number 2019/0638, approved February 19, 2020).

### Cell culture

SK-ES-1 ES cells obtained from the American Type Cell Collection (ATCC® HTB-86™) were verified for authenticity by Short Tandem Repeat (STR) and lack of contamination. Cells were cultured and incubated in 5% CO₂ at 37°C in RPMI-1640 Medium (Gibco™, Thermo Fisher Scientific, Waltham, USA), supplemented with 1% amphotericin B, 5% 10,000 Ul/mL penicillin 10 mg/ml streptomycin, and 10% fetal bovine serum (FBS, Gibco™, Thermo Fisher Scientific).

### Drug treatment

K252a, methyl (13S,14R,16R)-14-hydroxy-13-methyl-5-oxo-6,7,13,14,15,16-hexahydro-5H-13,16-epoxydiindolo[1,2,3-fg:3′,2′,1′-kl]pyrrolo[3,4-i][1,6]benzodiazocine-14-carboxylate, (Sigma Aldrich, St. Louis, USA) and the selective IGF1R inhibitor NVP-ADW742 (Sigma Aldrich) were dissolved in dimethyl sulfoxide (DMSO, Sigma Aldrich). K252a was dissolved in saline solution (NaCl 0.9%) at the concentration of 2% DMSO in 98% saline for the treatment of mice at a dose of 0.5 mg/kg. Control mice were given vehicle (2% DMSO in 98% saline). For *in vitro* assays, cells were treated with the inhibitory concentration of 10% (IC10) of K252a (10 nM), NVP-ADW742 (196.6 nM) or K252a plus NVP-ADW742. Control cells were exposed to vehicle (DMSO 0.2%) in the culture medium.

### Xenograft assay

Six-week-old male nude mice (BALB/c nu/nu) from our own institutional breeding colony were used for the *in vivo* experiments. Mice were randomly distributed into two groups, K252a treatment and control. A cell line-derived tumor xenograft (CDX) model was used to induce tumors. Animals were inoculated subcutaneously (s.c..) on the right dorsal flank with a suspension of SK-ES-1 cells (3 × 10⁶). After the tumors reached 80 mm³-100mm³, the mice received intraperitoneal (i.p.) injections with K252a (0.5 mg/kg/day) or the same volume of vehicle (2% DMSO in 98% saline) once daily during 18 days. Body weight (g), tumor width (W) and tumor length (L) were measured every three days until treatment day 18, and tumor volume (mm3) was determined by the formula (W² x L)/2. At the end of the treatment, the mice were euthanized under deep isoflurane anesthesia and the blood and tumors were collected for biochemical and histological analysis.

### Biochemical markers

Heart blood was collected via puncture pre-euthanasia and serum was separated (at 3,000 rpm for 10 min) and stored at −80 °C. Alanine aminotransferase (ALT), aspartate aminotransferase (AST), creatinine and urea serum levels were evaluated by closed-circuit photometric assays (Cobas® c702, F. Hoffmann-La Roche Ltd., Basel, Switzerland).

### Immunohistochemistry on tissue microarrays (TMA)

Tumors (*n* = 7 per group, removed immediately after the mouse’s death) were fixed in 10% formalin to be included in paraffin blocks. Sections of three micrometers were used for the production of slides stained with hematoxylin-eosin (HE) as they served to evaluate the morphology and an area for the fabrication of the tissue microarray (TMA). A 2 mm block of TMA was used for the study, where duplicates were posted as the source area. Ten antibodies were used: TrkA polyclonal antibody (Product # PA5-90581), TrkB polyclonal antibody (Product # PA5-88293), phospho-TrkA (Tyr791) polyclonal antibody (Product # PA5-105456), TrkC polyclonal antibody pTyr516 (Product # PA5-39755), phospho-IGF1R polyclonal antibody (Tyr1165, Tyr1166) (Product # PA5-104772), PI3K p85 alpha polyclonal antibody (Product # PA5-29220), phospho-PI3 kinase p85 alpha pTyr607 polyclonal antibody (Product # PA5-38905), phospho-TrkB polyclonal antibody pTyr515 (Product # PA5-38076), TrkC polyclonal antibody (Product # PA5-79766), TrkC polyclonal antibody (Product # PA5-79766), IGF1R (CD221) polyclonal antibody (Product # PA5-79445), and an anti-rabbit secondary antibody. The immunohistochemistry (IHC) protocol was carried out according to the specifications of each data sheet. The IHC assessment was scored semi-quantitatively according to intensity and distribution.

### Cell viability

SK-ES-1 ES cells were seeded at a density of 4 × 103 cells per well in a complete medium into 96-well plates (Kasvi, Pinhais, Brazil). After 24 h, cells were treated with K252a, NVP-ADW742, or the two drugs combined, for 72 h and the medium was then removed, cells were washed with phosphate-buffered saline (PBS) and detached with 0.25% trypsin solution (Gibco™, Thermo Fisher Scientific). For cell viability analysis, the cell suspension was homogenized with trypan blue (BioSumos, Porto Alegre, Brazil) and counted in a hemocytometer. The mean of three independent experiments for each dose was used to calculate dose-response and combination index values.

### Drug interactions

The combination index between the effects of K252a and NVP-ADW742 was calculated using the SinergyFinder/R version 4.1.3 package [57], with cell viability of SK-ES-1 ES cells as input using the ZIP (Zero Interaction Potency) method.

### Gene expression and survival analysis in ES patients

The gene expression dataset used for the survival analysis were obtained from the Gene Expression Omnibus (GEO) datasets accession numbers GSE63155 and GSE63156. The datasets GSE63155 and GSE63156 comprise expression data (Affymetrix Human Exon 1.0 ST Array) from primary ES tumor biopsies obtained from 46 patients from the Children’s Oncology Group (COG) and 39 patients from the EuroEwing collaborative group respectively [58]. Microarray raw data (GSE63155 and GSE63156, Affymetrix Human Exon 1.0 ST Array, GPL5175) normalization (Robust Multichip Average, RMA method) and quality control were performed using the oligo Bioconductor/R version 4.3.1). Overall survival (OS) analysis for ES patients was done by Kaplan-Meier curves using the ‘Survival’ and ‘Survminer’ packages, to evaluate the effects of expression of each gene of interest namely the NTRK1, NTRK2, NTRK3, IGF1R, PIK3CA (which encodes the p110α protein, the catalytic subunit of the PI3K), and PIK3R1 (which encodes a regulatory subunit of PI3K).

### Statistical analysis

The Shapiro-Wilk test and histograms were used to verify the distribution of variables. An independent t-test was used for comparisons between groups, assuming a parametric distribution. Repeated measurements (body weight and tumor volume) were assessed by the Generalized Estimating Equation (GEE). The GEE model was modeled to identify statistical differences between treatment groups in mice (K252a versus controls), as well as tumor volume between treatment days and the interaction between both factors. In the presence of a significant interaction indicated by *p* < 0.05, Bonferroni’s correction was used for post-hoc multiple comparisons. The ordinal histology variables were analyzed using the Spearman’s correlation method. All analyses were performed using the SPSS 20.0 software (IBM, Armonk, USA), assuming significance level at *p* < 0.05. Data are presented as mean and + standard error of the mean (S.E.M.).

## Supporting information

Supplementary information dos-Santos et al

## DATA AVAILABILITY

The ES tumor datasets GSE63155 and GSE63156, 28 used for the analyses of associations between gene expression and patient survival presented in this study, are available at https://www.ncbi.nlm.nih.gov/geo/query/acc.cgi?acc=GSE63155 and https://www.ncbi.nlm.nih.gov/geo/query/acc.cgi?acc=GSE63156, respectively. Other data presented in this study may be obtained from the corresponding author upon request.

## SUPPLEMENTARY MATERIALS

The following supporting information is provided: Supplementary Figure S1; Supplementary Figure S2; Supplementary Figure S3.

## AUTHOR CONTRIBUTIONS

Conceptualization, B.A.S. and C.B.F.; methodology, B.A.S., N.H.F., L.F., M.D., J.M.C., L.F.R.R., and M.C.J.; validation, M.A.C.F., M.C.J., and C.B.F.; formal analysis, B.A.S., C.B.F., M.D., and J.M.C.; investigation, B.A.S., N.H.F., L.F., M.D., J.M.C., L.F.R.R., and M.C.J.; resources, M.A.C.F., A.T.B., A.L.B., M.C.J., R.R., and C.B.F.; data curation, B.A.S., C.B.F., M.D., and J.M.C.; writing—original draft preparation, B.A.S., C.B.F. M.D., and R.R.; writing—review and editing, B.A.S., N.H.F., L.F., M.D., J.M.C., M.A.C.F., L.G., A.T.B., A.L.B., L.F.R.R. M.C.J., R.R., C.B.F.; supervision, M.A.C.F., L.G., A.T.B., A.L.B., R.R., C.B.F.; project administration, C.B.F.; funding acquisition, A.T.B., A.L.B., M.C.J., R.R., C.B.F. All authors have read and agreed to the published version of the manuscript.

## INSTITUTIONAL REVIEW BOARD STATEMENT

All experimental procedures involving mice were conducted in accordance with the guidelines established by the National Council for Animal Control and Experimentation (CONCEA, Ministry of Science, Technology and Innovation, Brazil), as well as with Animal Research: Reporting of In Vivo Experiments (ARRIVE) guidelines. All protocols used in this study were approved by the institutional Animal Care and Use Committee (CEUA, Hospital de Clínicas de Porto Alegre-HCPA), under project number 2019/0638, approved February 19, 2020.

