## Supplementary information dos-Santos et al for "Delayed growth of SK-ES-1 Ewing sarcoma tumors associated with reduced expression and activity of Trk receptors, PI3K, and IGF1R"

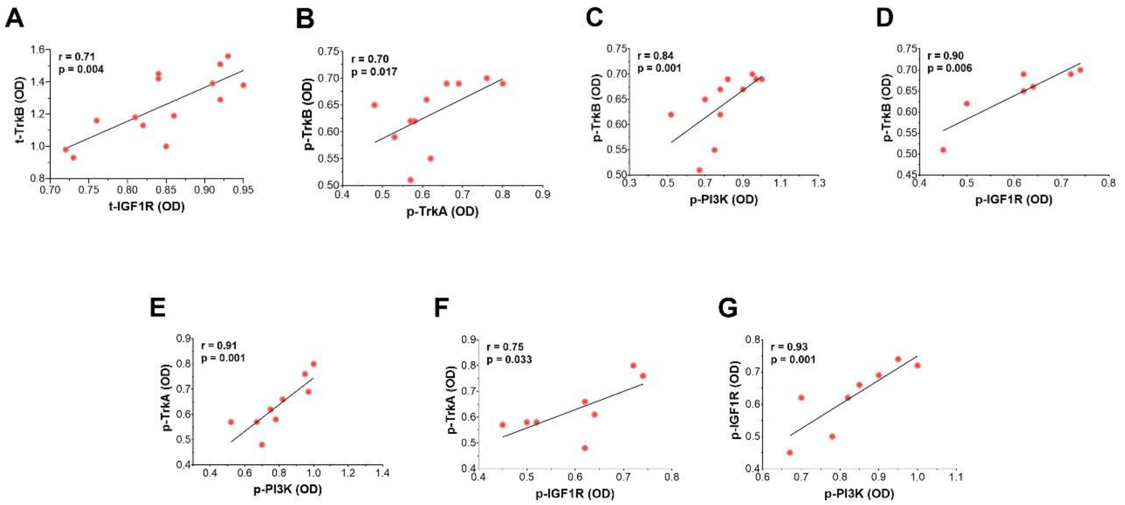


**Supplementary Figure S1.** Correlation analysis for the expression between (**A**) t-TrkB and t-IGF1R, (**B**) p-TrkB and p-TrkA, (**C**) p-TrkB and p-PI3K, (**D**) p-TrkB and p-IGF1R, (**E**) p-TrkA and p-PI3K, (**F**) p-TrkA and p-IGF1R, and (**G**) p-IGF1R and p-PI3K in ES in ES tumors from both the control and K252a-treated groups, selected on the basis of quality of immunhistochemical staining; n = 7-14.

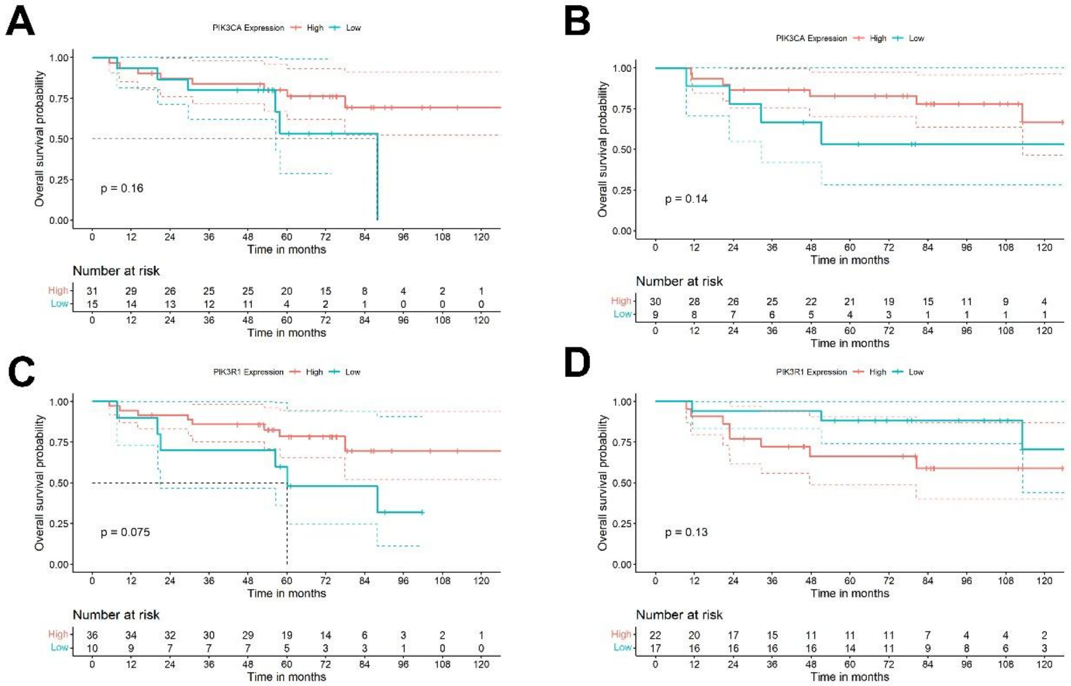


**Supplementary Figure S2.** Lack of significant associations between expression of genes encoding subunits of PI3K and prognosis assessed by OS in patients with ES. Datasets

GSE63155 and GSE63156, comprising expression data from 46 primary ES tumors biopsies from COG and 39 tumors from the EuroEwing collaborative group respectively [28] were used (**A**) *PIK3CA*, COG; (**B**) *PIK3CA*, EuroEwing, (**C**) *PIK3R1*, COG; (**D**) *PIK3R1*, EuroEwing; *p* values are indicated in each panel.

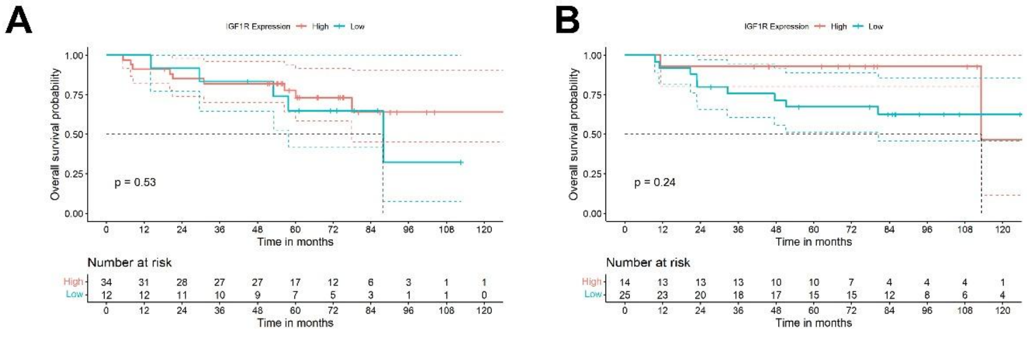


**Supplementary Figure S3.** Lack of significant association between *IGF1R* gene expression and prognosis assessed by OS in patients with ES. Datasets GSE63155 and GSE63156, comprising expression data from (**A**) 46 primary ES tumors biopsies from COG and (**B**) 39 tumors from the EuroEwing collaborative group respectively [58] were used; *p* values are indicated in each panel.
